# Mixed support for the idea that lower elevation animals are better competitors than their upper elevation relatives

**DOI:** 10.1101/634964

**Authors:** Benjamin G. Freeman

## Abstract

What factors set species’ range edges? One general hypothesis, often attributed to Darwin and MacArthur, is that interspecific competition prevents species from inhabiting the warmest portions along geographic gradients (i.e., low latitudes or low elevations). A prediction arising from the Darwin-MacArthur hypothesis is that lower elevation species are better competitors than are related upper elevation species. I tested this prediction by conducting a meta-analysis of studies that have measured behavioral competition between related species along elevational gradients. I found that (1) interspecific aggression appears to be a reliable indicator of interspecific competition; (2) as predicted, lower elevation species showed stronger interspecific aggression, but only for tropical species-pairs tested with playback experiments (nearly all songbirds); (3) for a broader range of taxa where authors directly observed behavioral interactions, *upper* elevation species showed stronger interspecific aggression; and (4) in general, larger species showed greater interspecific aggression. One potential explanation for why upper elevation species often show more interspecific aggression is that they tend to be larger (Bergmann’s rule; larger body sizes in colder environments). Supporting this possibility, tropical species tested with playback experiments, which do not follow Bergmann’s rule, were the only group that matched predictions arising from the Darwin-MacArthur hypothesis. Hence, available evidence does not consistently support the longstanding hypothesis that relative range position predicts the outcome of interspecific competition. Instead, body size is a better predictor of behavioral competition. Last, I consider these results in the context of the hypothesis that behavioral interactions may impact rates of upslope range shifts associated with recent warming.

---

Since Darwin, biologists have proposed that species interactions set species’ warm range edges, while climate sets species’ cold range edges (Darwin, 1859; MacArthur, 1972). A geographic prediction arising from this “Darwin-MacArthur” hypothesis is that species interactions—often thought to be competition—limit species’ ranges at low latitudes or low elevations (Louthan *et al.*, 2015). While some comparative studies have revealed patterns consistent with this prediction (Sunday *et al.*, 2012; Hargreaves *et al.*, 2014), others have shown opposite patterns (Cahill *et al.*, 2014; Freeman *et al.*, 2018). Therefore, whether competition (or other interactions) limit species’ warm range edges in nature remains an open question. One useful way to evaluate predictions arising from the Darwin-MacArthur hypothesis is to examine interactions between related species that live in different latitudinal or elevational zones. Here, I take this approach, focusing on competition along mountain slopes: I conducted a meta-analysis to test the prediction that species living at low elevations are better competitors than related species living at high elevations.

In this study, I focus on the behavioral component of competition (interference competition). Interspecific interference competition is often mediated by interspecific aggression (Schoener, 1983; Dhondt, 2011; Grether *et al.*, 2017), such that a better competitor will tend to exhibit more interspecific aggression in an interaction with a related species. If so, then the prediction arising from the Darwin-MacArthur hypothesis is that lower elevation species should be more aggressive to upper elevation relatives than vice versa. To test this prediction, I compiled a dataset of studies that have measured interspecific aggression between species-pairs where the two species live primarily in different elevational zones. A key assumption of my study is that interspecific aggression is a valid proxy for interspecific competition; I find support for this assumption when analyzing the subset of studies that measured both interspecific aggression and interspecific competition (Table 1, also see Results).

**Table 1.**
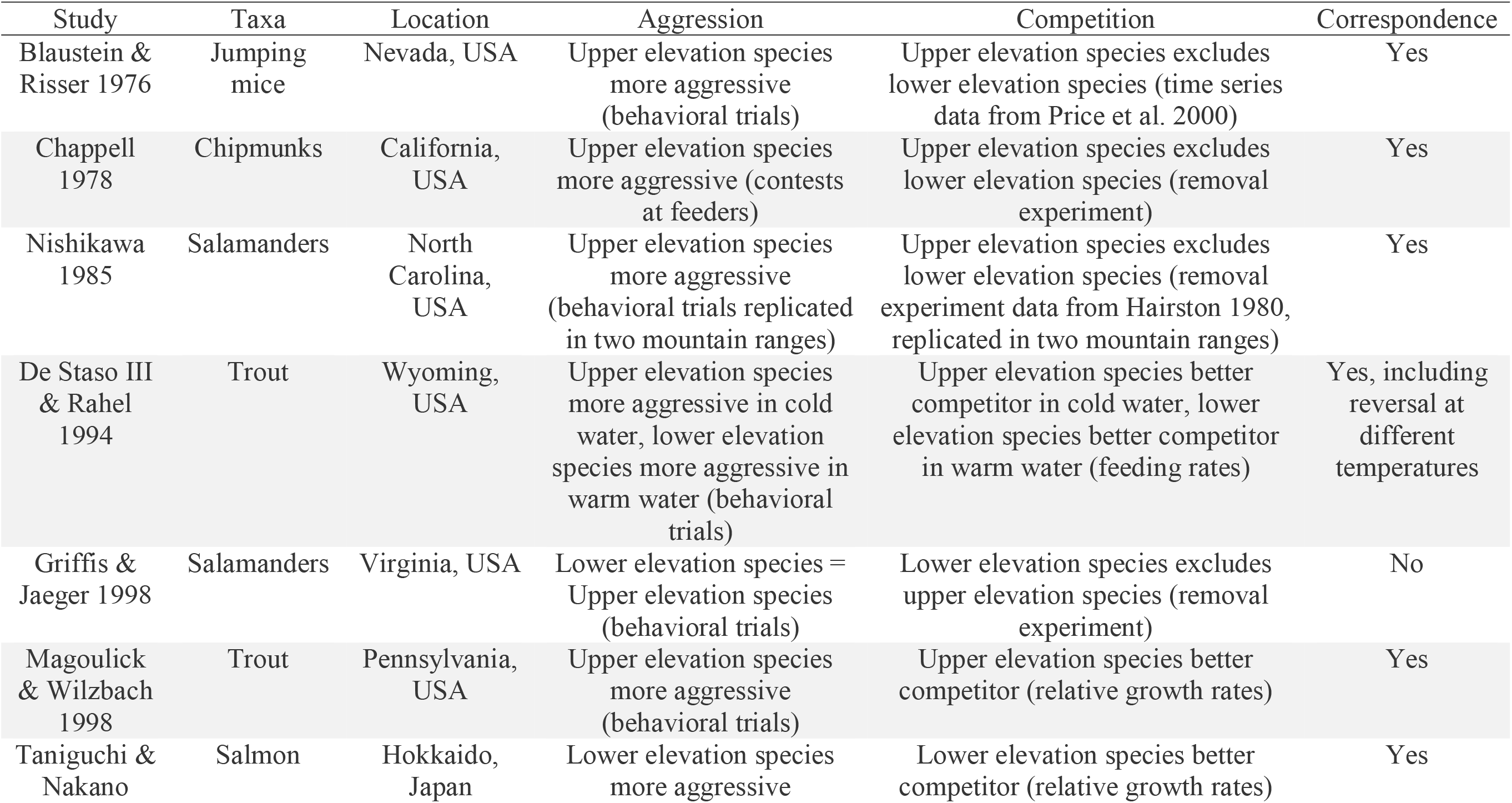
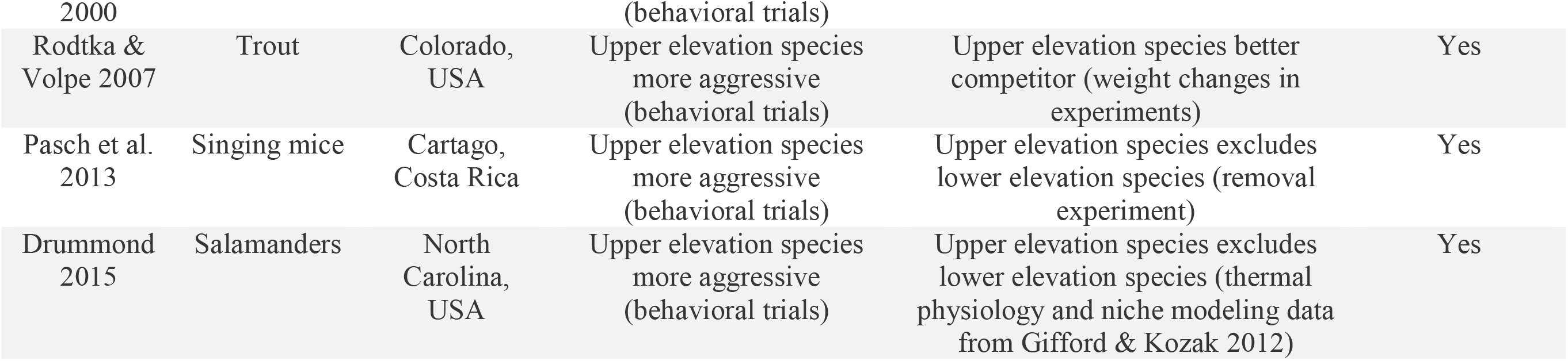
Details for 10 studies that measured both interspecific aggression and interspecific competition between lower vs. upper elevation species. The methods used to infer aggression and competition are given as parentheticals. The key question is whether interspecific aggression indicates interspecific competition. If the more aggressive species is also the better competitor, correspondence = “Yes”. Studies are arranged by year of publication.

A further motivation for testing general patterns of interspecific aggression as a function of elevational position is that behavioral interactions have been hypothesized to influence species’ geographic responses to climate change. The idea is that while we might generally expect species to shift upslope as a consequence of warming temperatures (as indeed is happening in nature; Chen *et al.*, 2011; Lenoir & Svenning, 2015; Freeman *et al.*, 2018), behavioral interactions could lead to faster or slower shifts over short time scales. That is, more aggressive lower elevation species might be able to rapidly colonize higher elevations, in the process “pushing” upper elevation relatives to ever higher elevations at rates faster than expected based solely on temperature changes; alternately, more aggressive upper elevation species might be able to avoid range contractions at their lower elevation (warm) range edge, persisting as “kings of the mountain”, at least in the short term (Jankowski *et al.*, 2010). The possibility that behavioral interactions influence rates of warming-associated upslope shifts has yet to be tested. Still, it is uncertain which of these situations—lower elevation species more aggressive (consistent with the Darwin-MacArthur hypothesis) vs. upper elevation species more aggressive—predominates in the real world. Hence, this meta-analysis both tests a longstanding hypothesis in ecology, and also provides information that is potentially relevant to understanding and predicting contemporary range shifts along mountain slopes.

## Methods

I searched the literature to find studies that measured interspecific aggression between species that live in different elevational zones along mountain slopes. I conducted a Web of Science search on 18 April 2019 with the keywords “behav*” OR “aggress*” AND “elevation*” OR “altitu*” AND “compet*. This search returned 561 studies. I retained studies that met the following two criteria: (1) They measured aggressive interactions between two closely related species (typically congeners), and (2) The two species in question inhabit roughly parapatric elevational zones during the breeding season (or all year long), with one species predominately living at lower elevations and the other at higher elevations. Disappointingly, this Web of Science search failed to return several older relevant papers that are routinely cited within this literature (e.g., Brown, 1971; Heller, 1971). Because my overall goal was to compile a comprehensive dataset, I located additional appropriate studies by (1) inspecting citations within papers identified by the Web of Science search, and (2) following citation webs. The final dataset included 57 estimates of interspecific aggression for 47 unique species-pairs from 34 studies. While the majority of estimates came from the temperate zone (absolute latitude > 23.4; N = 36), the tropics (absolute latitude < 23.4; N = 21) were also well represented (Figure 1). Taxonomically, the dataset consists of estimates for birds (N = 28), mammals (N = 12, mostly chipmunks from western North America), amphibians (N = 7, all *Plethodon* salamanders from the Appalachian Mountains in eastern North America), fishes (N = 8, all salmonids from temperate regions), reptiles (N = 1) and insects (N = 1).

**Figure 1.**
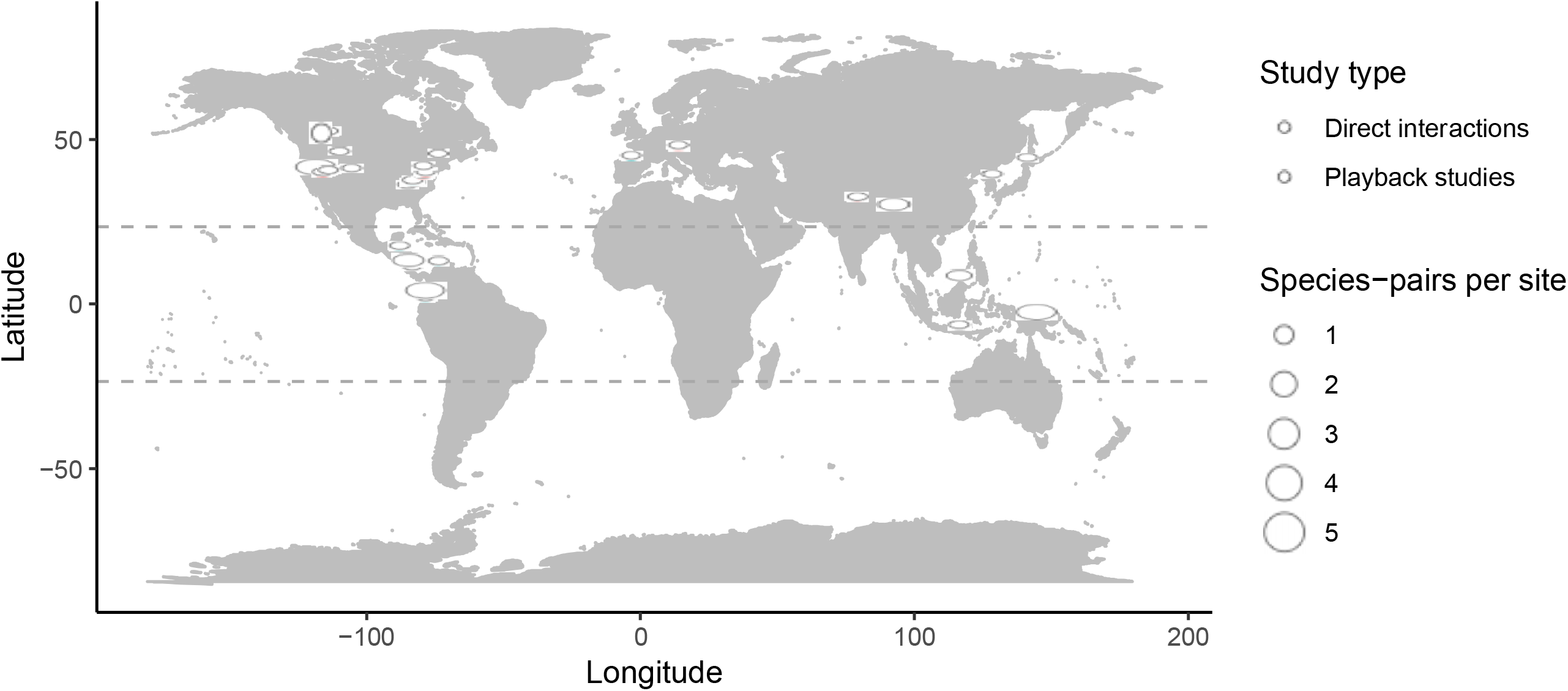
Map of studies that measured interspecific aggression between lower vs. upper elevation species. Studies that directly observed aggressive interactions are shown in pink; those that used playback to simulate interactions are shown in blue. Many studies report data for multiple species-pairs from the same site, illustrated by the size of the circle. The Tropics of Cancer and Capricorn (at 23.4° N and S, respectively) delimit the tropics, and are illustrated with dashed lines. This map was made using the package “ggmap” (Kahle & Wickham, 2013)

Upon examining relevant studies, I found a fundamental distinction between instances where authors directly observed interspecific aggression (N = 35), and those where authors measured aggressive behaviors in response to the simulated presence of a heterospecific (N = 22). Direct interaction studies measured aggression during contests in laboratory behavioral trials or at food sources placed in the environment (e.g., feeders). In contrast, simulated interaction studies measured aggression in response to song playback experiments, and were nearly always (21 out of 22) conducted on songbirds. Importantly, direct interaction vs. simulated playback studies (hereafter “direct” and “playback”) reported different metrics of interspecific aggression. I was unable to analyze these different metrics in a single meta-analytic model. Direct studies reported either the winner of aggressive contests or the number of aggressive behaviors exhibited during such contests. For these direct studies, I summarized interspecific aggression between the species-pair as a proportion—the proportion of contests won by the lower elevation species, or the proportion of total aggressive behaviors exhibited by the lower elevation species. Here, larger proportions indicate that the lower elevation species tends to win contests or exhibits more aggressive behaviors. This quantity can be directly used to test the prediction that lower elevation species are more aggressive to upper elevation relatives than vice versa. In contrast, playback studies reported aggression scores in response to a simulated heterospecific intruder (typically PC1 scores from a multivariate analysis of behavioral responses to playback, less often a univariate metric of aggression such as closest approach to the speaker). For these playback studies, I summarized interspecific aggression between the species-pair as an effect score—the difference in mean aggression scores between lower elevation and upper elevation species. Here, positive values represent cases where the lower elevation species showed more interspecific aggression than did the upper elevation species. Again, this difference can be directly used to test the prediction that lower elevation species are more aggressive to upper elevation relatives than vice versa. For all studies, when results were presented only in figures, I extracted data using WebPlotDigitizer (Rohatgi, 2017).

My principal aim in this study is to assess if lower elevation species show more interspecific aggression to upper elevation relatives than the reverse. I tested this idea by fitting mixed effect meta-analytic models using the “metafor” package (Viechtbauer, 2010) in R (R Development Core Team, 2017). I fit distinct models for direct and playback studies. These models weight individual estimates by the inverse of their squared standard errors, and incorporate the estimated variance among the study-specific effect sizes. Because my dataset included some species-pairs with multiple estimates of interspecific aggression (i.e., the same species-pair was measured in different studies), I included species-pair as a random effect in all models. For both direct and playback studies, I explored whether patterns differed between latitudinal zones or taxonomic groups by fitting secondary models that included either latitudinal zone or taxa as a moderator variable, and compared model fit using AIC. I did not fit a secondary model with taxonomic group for playback studies because nearly all playback studies were conducted on birds (21 out of 22 cases). The null expectation for the direct studies model is that the lower elevation species should win 50% of contests or exhibit 50% of observed aggressive behaviors. For playback studies, the null expectation is that lower and upper elevation species should have similar interspecific aggression scores, such that the true mean difference in aggression score is 0.

Last, I tested how body size was related to both interspecific aggression and elevational position. I extracted body size data (masses for birds and mammals, snout-vent-length for salamanders) from papers or, when not presented in papers, from reference volumes (Dunning, 2007; Wilman *et al.*, 2014) or personal communication with authors. I then used binomial tests to evaluate (1) if larger species showed more aggression to smaller species than vice versa (as expected, see Martin & Ghalambor, 2014), and (2) if upper elevation species tend to be larger than lower elevation species (as expected given Bergmann’s rule). For these analyses, I did not include playback studies where neither species showed interspecific aggression (defined as cases where response to song from the putative competitor was statistically indistinguishable from response to a negative control, a song from a totally unrelated species that is not expected to elicit any response). In sum I analyzed body mass for 33 unique species-pairs where at least one species showed interspecific aggression.

## Results

Interspecific aggression between related species along mountain slopes appears to be fairly common in nature. This is somewhat surprising, especially for playback studies. It is perhaps to be expected that individuals will behave aggressively when placed in a small laboratory arena with a single resource (i.e., most direct studies). But there is little expectation that simply broadcasting the song of a relative—a song that typically sounds obviously different from the focal species’ own song—should elicit an aggressive response. Nevertheless, the majority of species-pairs tested with playback studies in my dataset (15 out of 20) showed interspecific aggression, and interspecific aggression was as strong as intraspecific aggression in one-third of cases (7 out of 22; denominators differ because these inferences depend on experimental design, which varied among studies). Further, strong interspecific aggression appears to indicate competitive dominance, at least within this dataset. In the 10 studies that measured both interspecific aggression and interspecific competition, the more aggressive species was the better competitor in 9 out of 10 cases (binomial test; *p* = 0.021; Table 1).

I found mixed evidence that lower elevation species are more aggressive to upper elevation relatives than vice versa. For direct studies, the *upper* elevation species tended to win most contests or exhibit more aggressive behaviors in contests (Figure 2), the opposite of the predicted relationship. Competing models that included latitudinal zone or taxa as moderator variables provided poorer fits to the data (Δ AIC = 1.81 and 8.42, respectively). In contrast, in playback studies, lower elevation species did tend to exhibit more interspecific aggression (Figure 3), with an overall mean effect size that narrowly overlapped the null expectation of zero. A model that included latitudinal zone (tropical vs. temperate) provided a better fit to the data than a model without this moderator variable (Δ AIC = 1.66). In the model that included latitudinal zone, the estimate for tropical species-pairs was positive and did not overlap zero, indicating that the tendency for lower elevation species to be more aggressive in playback studies was associated with the tropics (Figure 3; the estimate for the subgroup of temperate zone species-pairs was approximately zero). While most studies conducted either direct observations or playback experiments, there were two studies that measured interspecific aggression using both direct observations and playback experiments, and these two studies both reported congruent results between the two methodologies (Pasch *et al.*, 2013; Barve & Dhondt, 2017).

**Figure 2.**
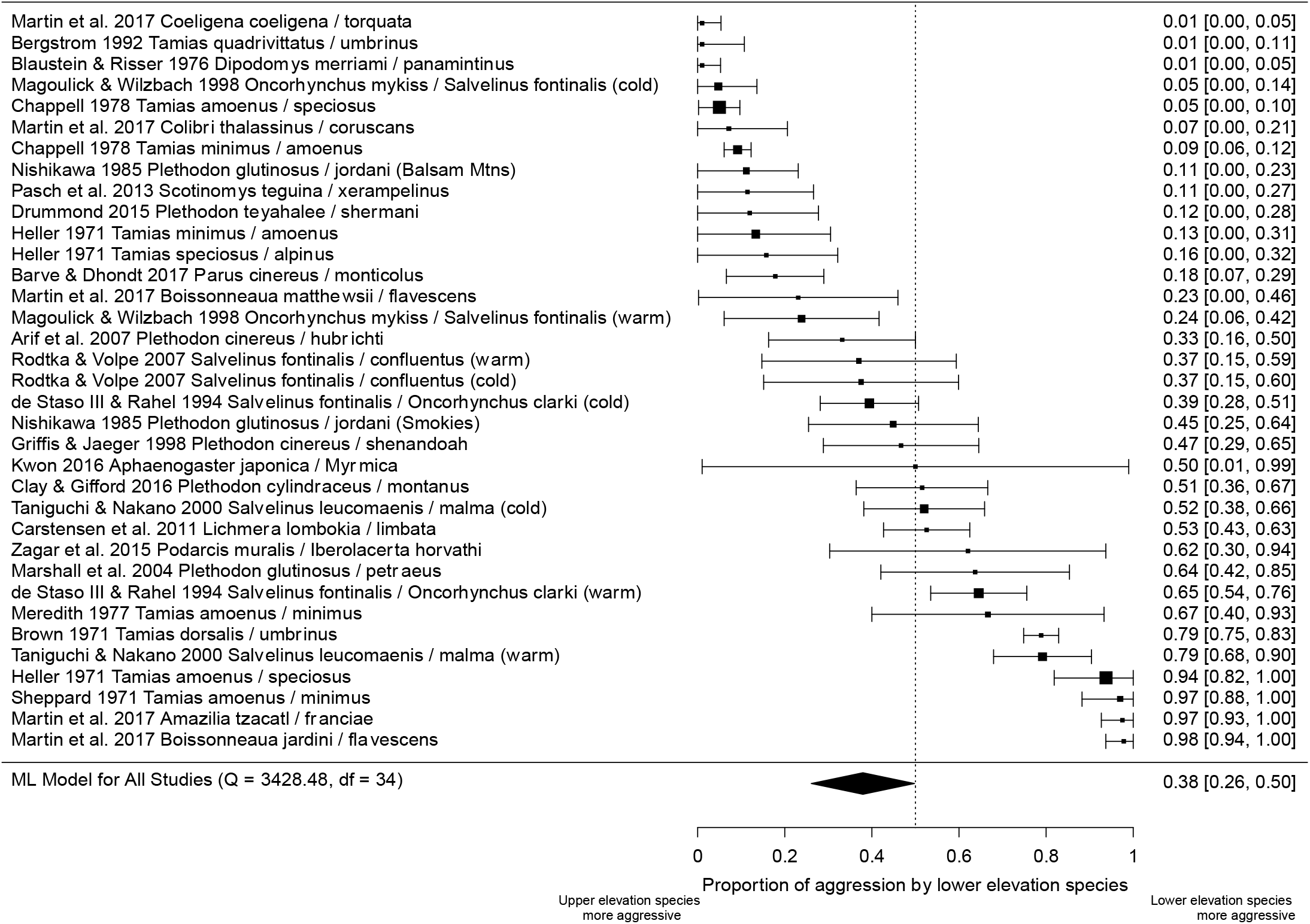
Results of studies that directly observed interspecific aggression (N = 36). The summary metric is the proportion of contests won, or aggressive behaviors exhibited, by the lower elevation species (± SE); this score would be 1 if the lower elevation species won all contests or exhibited 100% of all observed aggressive behaviors. Studies are identified by their author(s), year, and the scientific names of the species-pair.

**Figure 3.**
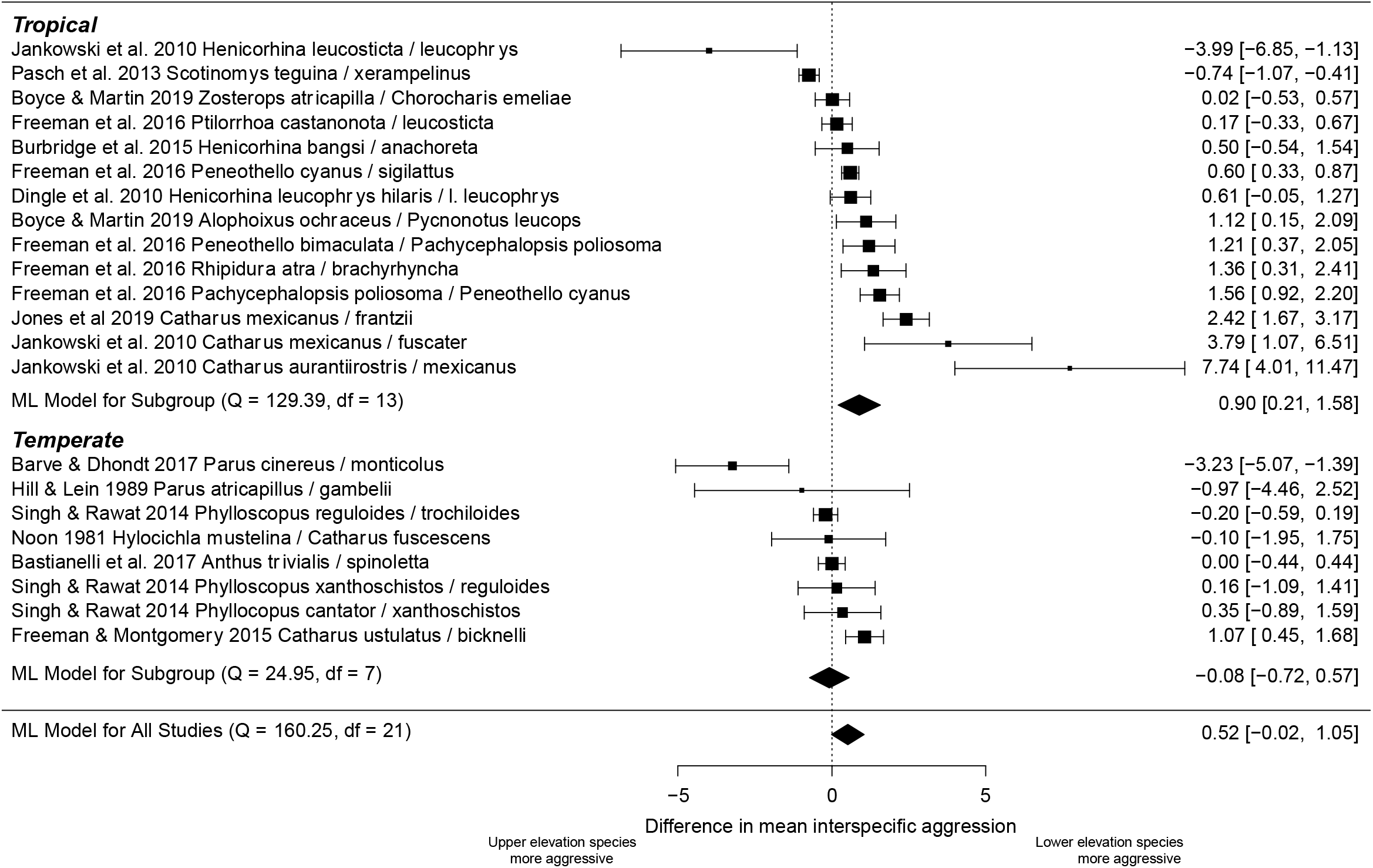
Results of studies that used playback experiments to measure interspecific aggression (N = 22), broken down into tropical and temperate zone subgroups. Effect sizes are calculated as the difference in mean interspecific aggression (± SE) between lower and upper elevation species, with positive values indicating cases where the lower elevation species exhibits stronger interspecific aggression than does the upper elevation species. Studies are identified by their author(s), year, and the scientific names of the species-pair.

I found evidence that body size drives observed patterns of interspecific aggression. Larger species tended to show stronger interspecific aggression (24 out of 33 species-pairs; binomial test; *p* = 0.014). However, overall, upper elevation species were not larger more often than by chance (18 out of 33 upper elevation species-pairs were larger; binomial test; *p* = 0.72). The association between size and elevational position differed somewhat between datasets (see Table 2), with upper elevation species tending to be larger in studies that directly observed behaviors (13 out of 21) but not in playback studies from the tropics (5 out of 11) [there were only three playback studies from the temperate zone that showed interspecific aggression; the upper elevation species was larger in two of these three cases].

**Table 2.** Patterns of body size, interspecific aggression, and elevational position for species-pairs from direct observation and playback studies. Relative size and relative interspecific aggression are coded as binary variables. Larger species tend to show more interspecific aggression than do smaller species (gray cells vs. white cells). In direct observation studies, but not playback studies (at least in the tropics), upper elevation species tend to be larger than lower elevation species (second column vs. first column). The numbers presented in this table sum to 35 (not 33) because two studies measured interspecific aggression using both direct observations and playback.

| Direct observation studies |  |  |
| --- | --- | --- |
|  | Lower = bigger | Upper = bigger |
| Lower = more aggressive | 5 | 2 |
| Upper = more aggressive | 3 | 11 |
| Playback studies (tropical) |  |  |
|  | Lower = bigger | Upper = bigger |
| Lower = more aggressive | 6 | 3 |
| Upper = more aggressive | 0 | 2 |
| Playback studies (temperate) |  |  |
|  | Lower = bigger | Upper = bigger |
| Lower = more aggressive | 1 | 1 |
| Upper = more aggressive | 0 | 1 |

## Discussion

I found mixed support for the prediction arising from the Darwin-MacArthur hypothesis that lower elevation species are more aggressive to their upper elevation relatives than vice versa. While lower elevation species showed stronger interspecific aggression in playback studies from the tropics, *upper* elevation species exhibited stronger interspecific aggression in studies that directly observed aggression, across a range of latitudes. Playback experiments were almost exclusively performed on forest-dwelling insectivorous birds, while direct observation studies investigated a much broader swath of biological diversity. Hence, I interpret this dataset as showing that insectivorous forest birds in the tropics show patterns consistent with the Darwin-MacArthur hypothesis, while patterns for other taxa, including for birds that are not tropical insectivores, are opposite to the prediction arising from the Darwin-MacArthur hypothesis. These mixed results indicate that the Darwin-MacArthur hypothesis is not a general explanation of aggressive interactions between related species along elevational gradients.

I next examine three reasons why my results do not consistently agree with predictions arising from the Darwin-MacArthur hypothesis, beginning with the possibility that my study is inappropriately designed. The most obvious caveat to my approach is that I attempt to study interspecific competition by analyzing patterns of interspecific aggression. That is, I assume interspecific aggression indicates competitive ability. This assumption need not hold—e.g., the most aggressive person lifting weights in your neighborhood gym may not necessarily win the local weightlifting competition. Nevertheless, this assumption does appear to be met in my dataset, as the more aggressive species was the better competitor in 9 out of 10 cases where researchers measured both variables (p = 0.021; Table 1). In addition, the subset of studies that measured interspecific competition are perhaps the most appropriate raw material for testing the Darwin-MacArthur hypothesis, but these “gold standard” studies fail to support predictions arising from this hypothesis. Instead, the *upper* elevation species was competitively superior to the lower elevation species in most cases where biologists measured competition (7 out of 9, see Table 1; note that competitive dominance flipped with temperature in one case).

Second, the evolution of larger body sizes at high elevations drivers could reverse expectations for patterns of interspecific aggression between low and high elevation species. That is, contra the Darwin-MacArthur hypothesis, we might expect *upper* elevation species to show more aggression towards lowland species than vice versa when two conditions are met— (1) larger species tend to win aggressive behavioral contests, and (2) species in colder high elevations are larger. These conditions likely apply broadly. The pattern that larger species tend to show more interspecific aggression than smaller species is strongly supported (Martin & Ghalambor, 2014), and Bergmann’s rule describes the well-known pattern that species in colder environments are larger. Indeed, larger species in my dataset exhibited more interspecific aggression in 73% of species-pairs (p = 0.014, see Table 2). In contrast, upper elevation taxa were larger in only just over half—55%—of species-pairs in my dataset (p = 0.73, see Table 2). Intriguingly, there is a rough correspondence between whether species-pairs show Bergmann’s rule body size clines and whether they show patterns consistent with predictions arising from the Darwin-MacArthur hypothesis. The only dataset that showed the predicted pattern of stronger interspecific aggression from lower elevation species was for tropical species-pairs (nearly all birds) from playback experiments, and there was no trend for upper elevation species to be larger within this subset (see Table 2). This is consistent with previous research showing that tropical birds do not generally follow Bergmann’s rule (Freeman, 2017). In contrast, upper elevation species did tend to be larger than lower elevation species in taxa in the direct observation dataset (where upper elevation species showed stronger interspecific aggression) and also for the small number of temperate zone species (Table 2). Hence, one possibility is that the evolution of larger body sizes at higher, colder elevations, at least in non-tropical birds, in conjunction with an advantage for larger body size in interspecific interference competition, may explain why the Darwin-MacArthur hypothesis does not generally apply to global patterns of interspecific aggression along mountain slopes. Further research with larger sample sizes is necessary to test this possibility.

Third, the simple logic of the Darwin-MacArthur hypothesis may require further refinement (see also Louthan *et al.*, 2015). Darwin’s proposal was that harsh climates prevent species from colonizing polar regions or high elevations, but that climate alone is unlikely to prevent species from colonizing the tropics or low elevations (Darwin, 1859). Instead, Darwin thought that tropical or low elevation species would have greater competitive ability relative to related temperate zone or high elevation species. This greater competitive ability would then prevent temperate zone/high elevation species from expanding at their warm range edge. However, if this basic scenario holds, with species’ warm range edges limited primarily by competition (and not by abiotic harshness), then selection on individuals’ competitive abilities would be strong at their warm range edge. If so, we might actually expect the evolution of increased competitive ability at species’ *warm* range edges. One prediction of this idea might be that shared range edges between lower vs. upper elevation species are located not where the lower elevation species meets unfavorable climatic conditions, but instead where the competitive balance tips between species adapted to different abiotic environments. In this case we might expect flips in competitive abilities along the elevational gradient, with the lower elevation species a better competitor at lower, warmer elevations, and the upper elevation species a better competitor at upper, cooler elevations (condition-dependent competition; see examples in Woodward, 1975; Taniguchi & Nakano, 2000; De Staso & Rahel, 2004; Altshuler, 2006). In general, investigating how selection on competitive ability varies across species’ ranges is likely to offer fresh insights for when competition sets species’ range edges.

### Behavioral interactions and range shifts

There have been widespread calls to incorporate species interactions when attempting to understand or predict species’ geographic responses to climate change (e.g., Araújo & Luoto, 2007; Alexander *et al.*, 2015). Behavioral interactions have the potential to be important for range limits. Indeed, case studies show that interspecific aggression appears to promote recent dramatic range expansions of native taxa (Duckworth & Badyaev, 2007; Wiens *et al.*, 2014). Extended to climate-associated range shifts, the idea is that aggressive lower elevation species could rapidly colonize elevations beyond their upper limit, “pushing” upper elevation relatives upslope, or that aggressive upper elevation species could hold steady at their lower limit as “kings of the mountain” despite warming. However, whether behavioral competition influences range shifts associated with climate change remains an open question. Consistent with a potential role for behavioral interactions in driving recent upslope shifts, I and colleagues reported that three aggressive lower elevation species of tropical birds have expanded their ranges upslope associated with recent warming while two non-aggressive lower elevation species have failed to expand (Freeman *et al.*, 2016). But another empirical example shows the reverse case—in the last century, a high elevation chipmunk *(Tamias speciosus)* has dramatically shifted upslope in the Sierra Nevadas in California (Moritz *et al.*, 2008) despite being a better behavioral competitor than a related lower elevation chipmunk (Heller, 1971). Clearly, many more studies are needed before we can confidently state that behavioral interactions between lower vs. upper elevation species do (or do not) predict species’ distributional responses to recent climate change.

### Conclusions

Probing the drivers of species’ range edges—explaining why a species lives *here* but not *there*—is a fundamental goal of ecology. More recently, this basic research has been reborn as an applied question, as ecologists are tasked with predicting where species will live in a warmer and climatically different future. The idea that general rules govern range limits is alluring in both basic and applied contexts, and the proposal that species’ warm range edges are set by competition (here termed the Darwin-MacArthur hypothesis) is one such general rule. In this paper I find mixed support for the prediction arising from this hypothesis that lower elevation species are better competitors than their upper elevation relatives, at least for behavioral competition. The “glass half empty” response to this finding would be to jettison the Darwin-MacArthur hypothesis (conditioned on the relatively small sample size of this study and the particular prediction being tested). However, an alternative “glass half full” approach would be to add a wrinkle to the hypothesis—that lower elevation species may indeed tend to be better behavioral competitors than their upper elevation relatives, but only when they are similar in mass (i.e., when Bergmann’s rule does not apply). Testing which of these scenarios is better supported will require additional data. I was pleasantly surprised to find as many studies measuring behavioral interactions between species-pairs along mountain slopes as I did (57 estimates of interspecific aggression for 47 unique species-pairs from 34 studies), but more empirical data is needed to test whether Bergmann’s rule modifies the general expectation that competition sets warm range limits, or that behavioral interactions are important in driving climate-associated range shifts, at least in the short term. My hope is that this literature continues to expand, such that someone revisiting this topic in a decade’s time will be able to provide firm answers to these and related questions.

## Acknowledgements

This research would not be possible without the many dedicated researchers who have measured and published behavioral interactions along mountain slopes—thank you and keep up the good work. I was supported by postdoctoral fellowships from the Biodiversity Research Centre and Banting Canada (#379958). None of my funders had any influence on the content of the submitted manuscript, and none of my funders required approval of the final manuscript to be published. Comments from the Schluter lab group, Locke Rowe and Ralf Yorque improved this manuscript.

